# mixOmics: an R package for ‘omics feature selection and multiple data integration

**DOI:** 10.1101/108597

**Authors:** Florian Rohart, Benoît Gautier, Amrit Singh, Kim-Anh Lê Cao

## Abstract

The advent of high throughput technologies has led to a wealth of publicly available ‘omics data coming from different sources, such as transcriptomics, proteomics, metabolomics. Combining such large-scale biological data sets can lead to the discovery of important biological insights, provided that relevant information can be extracted in a holistic manner. Current statistical approaches have been focusing on identifying small subsets of molecules (a ‘molecular signature’) to explain or predict biological conditions, but mainly for a single type of ‘omics. In addition, commonly used methods are univariate and consider each biological feature independently.

We introduce `mixOmics`, an R package dedicated to the multivariate analysis of biological data sets with a specific focus on data exploration, dimension reduction and visualisation. By adopting a system biology approach, the toolkit provides a wide range of methods that statistically integrate several data sets at once to probe relationships between heterogeneous ‘omics data sets. Our recent methods extend Projection to Latent Structure (PLS) models for discriminant analysis, for data integration across multiple ‘omics data or across independent studies, and for the identification of molecular signatures. We illustrate our latest `mixOmics` integrative frameworks for the multivariate analyses of ‘omics data available from the package.

## I. Introduction

The advent of novel ‘omics technologies (*e.g.* transcriptomics for the study of transcripts, proteomics for proteins, metabolomics for metabolites, etc) has enabled new opportunities for biological and medical research discoveries. Commonly, each feature from each technology (transcripts, proteins, metabolites, etc) is analysed independently through univariate statistical methods including ANOVA, linear models or t-tests. However, such analysis ignores relationships between the different features and may miss crucial biological information. Indeed, biological features act in concert to modulate and influence biological systems and signalling pathways. Multivariate approaches, which model features as a set, can therefore provide a more insightful picture of a biological system, and complement the results obtained from univariate methods. Our package `mixOmics` proposes multivariate projection-based methodologies for ‘omics data analysis as those provide several attractive properties to the data analyst Lê Cao et al. (2017). Firstly, they are computationally efficient to handle large data sets, where the number of biological features (usually thousands) is much larger than the number of samples (usually less than 50). Secondly, they perform dimension reduction by projecting the data into a smaller subspace while capturing and highlighting the largest sources of variation from the data, resulting in powerful visualisation of the biological system under study. Lastly, their relaxed assumptions about data distribution make them highly flexible to answer topical questions across numerous biology-related fields Boulesteix and Strimmer (2007); Meng et al. (2016). mixOmics multivariate methods have been successfully applied to statistically integrate data sets generated from difference biological sources, and to identify biomarkers in ‘omics studies such as metabolomics, brain imaging and microbiome Labus et al. (2015); Cook et al. (2016); Guidi et al. (2016); Mahana et al. (2016); Ramanan et al. (2016); Rollero et al. (2016).

We introduce `mixOmics` in the context of *supervised analysis*, where the aims are to classify or discriminate sample groups, to identify the most discriminant subset of biological features, and to predict the class of new samples. We further extended our core method sparse Partial Least Square - Discriminant Analysis (sPLS-DA Lê Cao et al. (2011)) that was originally developed for the supervised analysis of one data set. Our two novel frameworks DIABLO and MINT focus on the integration of multiple data sets for different biological questions (Fig 1). DIABLO enables the integration of the same biological *N* samples measured on different ‘omics platforms (*N*-integration, Singh et al. (2016)), while MINT enables the integration of several independent data sets or studies measured on the same *P* predictors (*P*-integration, Rohart et al. (2017)). To date, very few statistical methods can perform *N*- and *P*-integration in a supervised context. For instance, *N*-integration is often performed by concatenating all the different ’omics data sets Liu et al. (2013), which ignores the heterogeneity between ‘omics platforms and mainly highlights one single type of ’omics. The other common type of *N*-integration is to combine the molecular signatures identified from separate analyses of each ‘omics Günther et al. (2012), which disregards the relationships between the different ‘omics functional levels. With *P*-integration, statistical methods are often sequentially combined to accommodate or correct for technical differences (‘batch effects’) among studies before classifying samples with a suitable classification method. Such sequential approaches are time consuming and are prone to overfitting when predicting the class of new samples Rohart et al. (2017). Our two frameworks model relationships between different types of ‘omics data (*N*-integration) or integrate independent ‘omics studies to increase sample size and statistical power (*P*-integration). Both frameworks aim at identifying biologically relevant and robust molecular signatures to suggest novel biological hypotheses.

**Figure 1:**
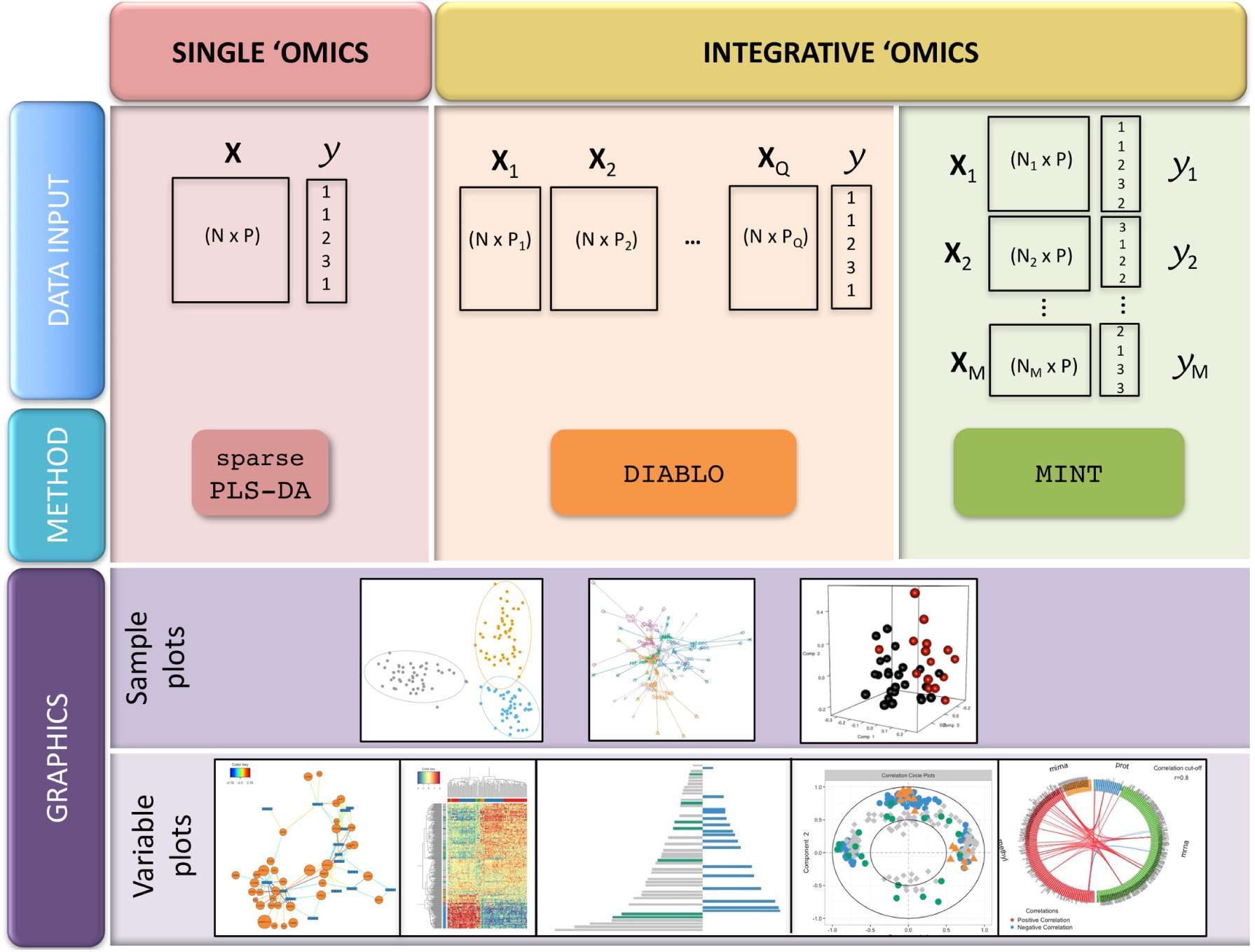
Overview of the mixOmics multivariate methods for single and integrative ‘omics supervised analyses. *X denote a predictor ‘omics data set, and y a categorical outcome response (e.g. healthy vs. sick). Integrative analyses include N-integration with **DIABLO** (the same N samples are measured on different ‘omics platforms), and P-integration with **MINT** (the same P ‘omics predictors are measured in several independent studies). Sample plots depicted here use the **mixOmics** functions (from left to right) **plotIndiv, plotArrow** and **plotIndiv** in 3D; variable plots use the **mixOmics** functions **network, cim, plotLoadings, plotVar** and **circosPlot**. The graphical output functions are detailed in Supporting Information.*

The present article first introduces the main functionalities of `mixOmics`, then presents our multivariate frameworks for the identification of molecular signatures in one and several data sets, and illustrates each framework in a case study available from the package. Reproducible Sweave code is provided for all analyses.

## Design and Implementation

`mixOmics` is a user-friendly R package dedicated to the exploration, mining, integration and visualisation of large data sets Lê Cao et al. (2017). It provides attractive functionalities such as (i) insightful visualisations with dimension reduction (Fig 1), (ii) identification of molecular signatures and (iii) improved usage with common calls to all visualisation and performance assessment methods (Supporting Information).

### Data input

Different types of biological data can be explored and integrated with mixOmics. Prior to the analysis, we assume the data sets have been normalised using appropriate techniques specific for the type of ‘omics technology platform. The methods can handle molecular features measured on a continuous scale (e.g. microarray, mass spectrometry-based proteomics and metabolomics) or sequenced-based count data (RNA-seq, 16S, shotgun metagenomics) that become “continuous” data after pre-processing and normalisation.

We denote *X* a data matrix of size *N* observations (rows) *× P* predictors (e.g. expression levels of *P* genes, in columns). The categorical outcome *y* (e.g. sick vs healthy) is expressed as a dummy indicator matrix *Y*, where each column represents one outcome category and each row indicates the class membership of each sample. Thus, *Y* is of size *N* observations (rows) *× K* categories outcome (columns), see example in Suppl.

While mixOmics methods can handle large data sets (several tens of thousands of predictors), we recommend pre-filtering the data to less than 10K predictors per data set, for example by using Median Absolute Deviation Teng et al. (2016) for RNA-seq data, by removing consistently low counts in microbiome data sets Arumugam et al. (2011); Lê Cao et al. (2016) or by removing near zero variance predictors. Such step aims to lessen the computational time during the parameter tuning process.

### Multivariate projection-based methods

mixOmics offers a wide range of multivariate dimension reduction techniques designed to each answer specific biological questions, via unsupervised or supervised analyses. The mixOmics functions are listed in Table 1. Unsupervised analyses methods includes Principal Component Analysis – based on NonLinear Iterative Partial Least Squares for missing values Wold (1975), Independent Component Analysis Yao et al. (2012), Partial Least Squares regression – PLS, also known as Projection to Latent Structures Wold (1966), multi-group PLS Eslami et al. (2013), regularised Canonical Correlation Analysis – rCCA González et al. (2008)) and regularised Generalised Canonical Correlation Analysis – RGCCA based on a PLS algorithm Tenenhaus and Tenenhaus (2011). Supervised analyses methods includes PLS-Discriminant Analysis – PLS-DA Nguyen and Rocke (2002b,a); Boulesteix (2004), GCC-DA Singh et al. (2016) and multi-group PLS-DA Rohart et al. (2017). In addition, mixOmics provides novel sparse variants that enable *feature selection*, the identification of key predictors (e.g. genes, proteins, metabolites) that constitute a *molecular signature*. Feature selection is performed via *l*^1^ regularisation (LASSO, Tibshirani (1996)), which is implemented into each method’s statistical criterion to be optimised. For supervised analyses, mixOmics provides functions to assist users with the choice of parameters necessary for the feature selection process (see ‘Choice of parameters’ Section) to discriminate the outcome of interest (*e.g.* healthy *vs.* sick, or tumour subtypes, etc.).

**Table 1:**
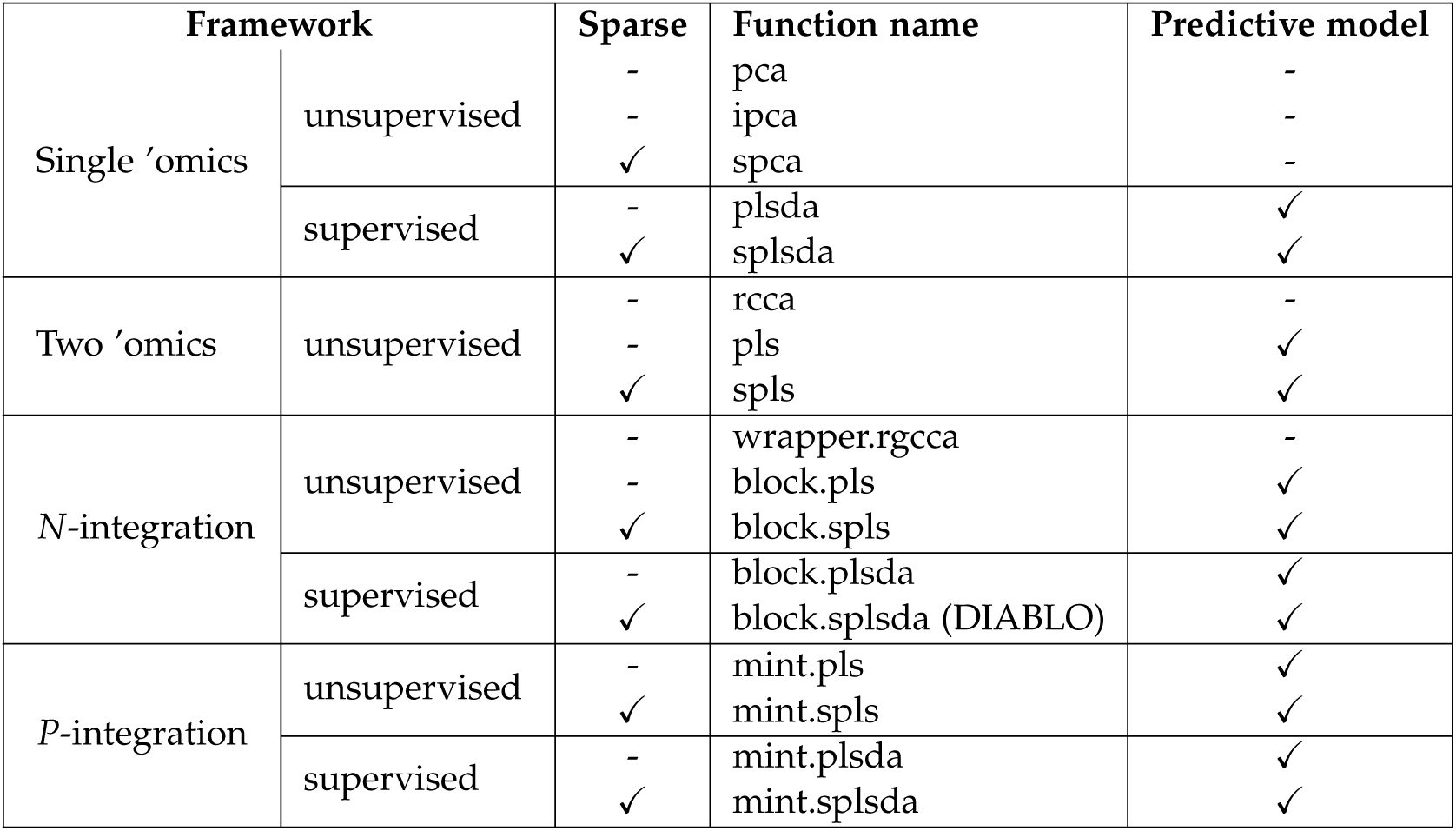
*Summary of multivariate projection-based methods available in mixOmics version 6.0.0 or above for different types of analysis frameworks. Note that our block.pls/plsda and sparse variants differ from the approaches from Wangen and Kowalski (1989); Westerhuis and Smilde (2001); Karaman et al. (2015); Kawaguchi and Yamashita (2017).*

All multivariate approaches listed in Table 1 are projection-based methods whereby samples are summarised by *H latent components* (*t*_1_, …, *t*_*H*_) that are defined as linear combinations of the original predictors. In the combinations (*t*_1_, …, *t*_*H*_), the weights of each of the predictors are indicated in the *loading vectors a*_1_, …, *a*_*H*_. For instance, for the data matrix *X* = (*X*^1^, …, *X*^*P*^) we define the first latent component as 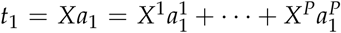. Therefore, to each loading vector *a*_*h*_ corresponds a latent component *t*_*h*_, and there are as many pairs (*t*_*h*_, *a*_*h*_) as the chosen *dimension H* in the multivariate model, *h* = 1, …, *H*, where *H << P*. The samples are thus projected into a smaller interpretable space spanned by the *H* latent components.

### Implementation

`mixOmics` is currently fully implemented in the R language and exports more than 30 functions to perform statistical analyses, tune the methods parameters and plot insightful visualisations. mixOmics mainly depends on the R base packages (parallel, methods, grDevices, graphics, stats, utils) and recommended packages (MASS, lattice), but also imports functions from other R packages (igraph, rgl, ellipse, corpcor, RColorBrewer, plyr, dplyr, tidyr, reshape2, ggplot2). In mixOmics, we provide generic R/S3 functions to assess the performance of the methods (predict, plot, print, perf, auroc, etc) and to visualise the results as depicted in Fig 1 (plotIndiv, plotArrow, plotVar, plotLoadings, etc), see Supporting Information for an exhaustive list.

Currently, seventeen multivariate projection-based methods are implemented in mixOmics to in-tegrate large biological data sets, amongst which twelve have similar names (mint).(block).(s)pls(da), see Table 1. To perform either *N*- or *P*-integration, we efficiently coded the functions as wrappers of a single main hidden and generic function that is based on our extension of the SGCCA algorithm Tenenhaus et al. (2014). The remaining five statistical methods are PCA, sparse PCA, IPCA, rCCA and rGCCA. Each statistical method implemented in mixOmics returns a list of essential outputs which are used in our S3 visualisation functions (Supporting Information).

`mixOmics` aims to provide insightful and user-friendly graphical outputs to interpret statistical and biological results, some of which (correlation circle plots, relevance networks, clustered image maps) were presented in details in González et al. (2012). The function calls are identical for all multivariate methods via the use of R/S3 functions, as we illustrate in the Results Section. mixOmics offers various visualisations, including sample plots and variable plots, which are based on latent component scores and loading vectors, respectively (Fig 1). Additional graphical outputs are available in mixOmics to illustrate classification performance of multivariate models using the generic function plot (see Supporting Information).

### Class prediction of new samples

PLS is traditionally a regression model where the response *Y* is a matrix of continuous data. To perform classification and prediction, supervised multivariate methods in mixOmics extend PLS by coding the categorical outcome factor as a dummy indicator matrix before being input into our PLS-based approaches. Considering an independent test set or a cross-validation set, the predict function calculates *predicted coordinates (scores*) and *predicted dummy variables* for each new observation, from which we obtain the final predicted class via the use of prediction distances (details in Supporting Information).

#### Prediction distances

For each new observation, we predict its coordinates on the set of *H* latent components, similarly to a multivariable multivariate model. These predicted coordinates, or scores, are then used to predict each of the *K* dummy variables. The predicted class of each new observation is derived by applying a distance to either the *H* predicted scores, or the *K* predicted dummy variables. We propose distances such as the ‘maximum distance’, ‘Mahalanobis distance’ and ‘Centroids distance’, which are detailed in Supporting Information. The maximum distance is applied to the predicted dummy variables and predicts the class category with the maximum dummy value. In single ‘omics analyses this distance achieves best accuracy when the predicted values are close to 0 or 1 Lê Cao et al. (2011). The ‘Mahalanobis distance’ and ‘Centroids distance’ are distances that are both applied to the predicted scores, and are based on the calculation of a centroid for each outcome category using the *H* latent components. These distances are appropriate for complex classification problems where samples should be considered in a multi-dimensional space spanned by the components. The predicted class of a new observation is the class for which the distance between its centroid and the *H* predicted scores is minimal, based on either the Euclidian distance (‘centroid distance’), or the ‘Mahalanobis distance’. The former assumes a spherical distribution around the centroid whereas the latter is more adapted for ellipsoidal distribution. In practice, we found that those distances, and especially the Mahalanobis distance, were more accurate than the maximum distance for N-integration. All distances consider the predictions built from all components of the model.

#### Visualisation of prediction area

To visualise the effect of the prediction distance, we propose a graphical output of the prediction area that overlays the sample plot (example in Fig 2 and more details in Supporting Information).

**Figure 2:**
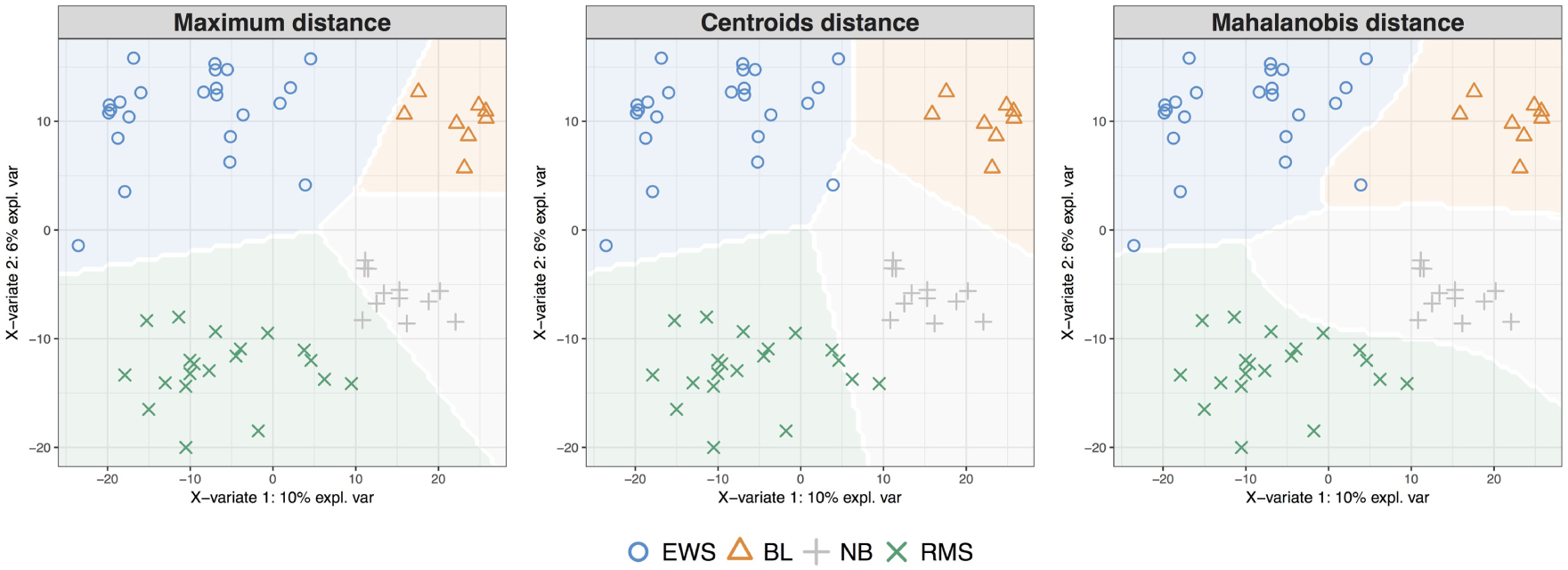
Prediction area visualisation. *on the Small Round Blue Cell Tumors data (SRBCT Khan et al. (2001)) data, described in the Results Section, with respect to the prediction distance. From left to right: ‘maximum distance’, ‘centroids distance’ and ‘Mahalanobis distance’. Sample prediction area plots from a PLS-DA model applied on a microarray data set with the expression levels of 2,308 genes on 63 samples. Samples are classified into four classes: Burkitt Lymphoma (BL), Ewing Sarcoma (EWS), Neuroblastoma (NB), and Rhabdomyosarcoma (RMS).*

#### Prediction for *N*-integration

For *N*-integration, we obtain a predicted class *per* ‘omics data set. The predictions are combined by majority vote (the class that has been predicted the most often across all data sets) or by weighted vote, where each ‘omics data set weight is defined as the correlation between the latent components associated to that particular data set and the outcome, from the training set. The final prediction is the class that obtains the highest weight across all ‘omics data sets. Therefore the weighted vote gives more importance to the ‘omics data set that is best correlated to the outcome and reduces the number of ties when an even number of data sets are discordant in the case of majority vote. Ties are indicated as NA in our outputs.

### Prediction for *P*-integration

In that specific case, the external test set can include samples from one of the independent studies used to fit the model, or samples from external studies, see Rohart et al. (2017) for more details.

### Choice of parameters for supervised analyses

For supervised analysis, mixOmics provides tools to choose the number of components *H* and the *l*^1^ penalty on each component for all sparse methods before the final multivariate model is built and the selected features are returned.

#### Parameter tuning using cross-validation

For all supervised models, the function tune implements repeated and stratified cross-validation (CV, see details in Supporting Information) to compare the performance of models constructed with different *l*^1^ penalties. Performance is measured via overall misclassification error rate and Balanced Error Rate (BER). BER is appropriate in case of an unbalanced number of samples per class as it calculates the average proportion of wrongly classified samples in each class, weighted by the number of samples in each class. Therefore, BER is less biased towards majority classes during the performance assessment. The choice of the parameters (described below) is made according to the best prediction accuracy, i.e. the lowest overall error rate or lowest BER.

#### Number of components

For all supervised methods, the tuning function outputs the optimal number of components that achieve the best performance based on the overall error rate or BER. The assessment is data-driven and similar to the process detailed in Rohart et al. (2016), where one-sided t-tests assess whether there is a gain in performance when adding components to the model. In practice (see some of our examples in the Results Section), we found that *K −* 1 components, where K is the number of classes, was sufficient to achieve the best classification performance Lê Cao et al. (2011); Shah et al. (2016). However, assessing the performance of a non sparse model with *K* to *K* + 2 components can be used to identify the optimal number of components, see Electronic Supporting files, and.

#### *l*^1^ penalty or the number of features to select

Contrary to other R packages implementing *l*^1^ penalisation methods (*e.g.* glmnet, Friedman et al. (2010), PMA, Witten et al. (2013)), mixOmics uses soft-thresholding to improve usability by replacing the *l*^1^ parameter by the number keepX of features to select on each dimension. The performance of the model is assessed for each value of keepX provided as a grid by the user from the first component to the *H*^*th*^ component, one component at a time. The grid needs to be carefully chosen to achieve a trade-off between resolution and computational time. Firstly, one should consider the minimum and maximum values of the selection size that can be handled practically for follow-up analyses (e.g. wet-lab experiments may require a small signature, gene ontology a large signature). Secondly, one should consider the computational aspect, as the tune function performs repeated cross-validation. For single ‘omics and P-integration analyses, a coarse tuning grid can be assessed first to evaluate the likely boundaries of the keepX values before setting a finer grid. For N-integration, as different combinations of keepX between the different ‘omics are assessed, a coarse grid is difficult to achieve as a preliminary step.

The tune function returns the set of keepX values that achieve the best predictive performance for all the components in the model. In case of ties, the lowest keepX value is returned to obtain a minimal molecular signature. The same grid of keepX values is used to tune each component; however for N-integration, different grids can be set for each data set. Examples of optimal keepX values returned by our functions are detailed in the Results section (see also , , and).

#### Special case for *P*- integration

For *P*-integration, we take advantage of the independence between studies. A Leave-One-Group-Out Cross Validation is performed where each study defines a subset that is left out once, as described in Rohart et al. (2017), which susbtantially improves computational time (see Supporting Information and for additional details).

### Evaluating the signature

#### Performance assessment

Once the optimal parameters have been chosen (number of components and number of variables to select), the final model is run on the whole data set *X*, and the performance of the final model in terms of classification error rate is estimated using the perf function and repeated CV. Additional evaluation outputs include the receiver operating characteristic (ROC) curves and Area Under the Curve (AUC) averaged over the cross-validation process using one-vs-all comparison if *K >* 2. AUC is a commonly used measure to evaluate a classifier discriminative ability. It incorporates the measures of sensitivity and specificity for every possible cut-off of the predicted dummy variables. However, as presented in Section ‘Prediction distances’, our PLS-based models rely on prediction distances, which can be seen as a determined optimal cut-off. Therefore, the ROC and AUC criteria may not be particularly insightful in relation to the performance evaluation of our supervised multivariate methods, but can complement the statistical analysis.

#### Stability

A by-product of the performance evaluation using perf is the record of the features that were selected across the (repeated) CV runs. The function perf outputs the feature stability per component to assess the reproducibility of the molecular signature (see example in Electronic Supporting).

#### Graphical outputs

Variable plots are useful to assess the correlation of the selected features within and between data sets. Correlation circle plots, clustered image maps, relevant networks are described in Supporting Information. A pyramid barplot displays the loading weights associated to each selected feature in increasing order of importance (from bottom to top), with colors indicating the sample group with the maximum or alternatively minimum average value (see Supporting Information).

### Computational aspects

The choice of the parameters via the tuning and the performance evaluation steps can be computationally demanding as the tune and perf function perform repeated cross-validation. Once the optimal parameters are chosen, the final multivariate models in mixOmics are however computationally very efficient to run.

The tuning can be particularly intensive for N-integration as we test all possible combination of subsets of variables to select. For large multi-‘omics data sets, the tuning will often require the use of a cluster, while a normal laptop might be sufficient for the single ‘omics and P-integration. To lessen the computational issue, the argument cpus in both tune and perf functions is included for parallel computing. Table 2 reports the computational time for the analyses illustrated in the Electronic Supporting Files , , and. The data analysed constitute a reduced set of features that are included in the package. Supplemental Information reports the computational time for large data sets analysed with mixOmics. For the latter case, we usually recommend to filter the data sets, as detailed in Section ‘Data input’ for a more tractable analysis.

**Table 2.** Example of computational time for the data sets presented in the Results section with a macbook pro 2013, 2.6GHz, 16Go Ram.

| Framework | Single 'omics<br>sPLS-DA |  | N-integration<br>DIABLO |  | P-integration<br>MINT |  |
| --- | --- | --- | --- | --- | --- | --- |
| Data | srbct |  | breast.tcga |  | stemcells |  |
| *N* | 63 |  | 150 |  | 125 |  |
| *P* | 2,308 |  | 200;184;142 |  | 400 |  |
| function | tune | perf | tune | perf | tune | perf |
| #fold CV (repeated) | 5(10) | 5(10) | 5(1) | 5(10) | LOGOCV | LOGOCV |
| ncomp | 5 | 3 | 2 | 2 | 2 | 2 |
| grid length per component | 39 | - | 13<sup>3</sup> | - | 100 | - |
| #cpu | 1 | 1 | 2 | 1 | 1 | 1 |
| run time | 2hrs | 27sec | 3hrs | 45sec | 35sec | 0.6sec |

## Results

We illustrate three supervised frameworks’ analyses performed with mixOmics using data available from the package. These data sets were reduced to fit the memory allocation storage allowed in R CRAN and the results presented are hence an illustration of the capabilities of our package, but do not necessarily provide insightful biological results.

### Single ‘omics supervised analyses with PLS-DA and sPLS-DA

We present the application of the single ‘omics multivariate methods PCA, PLS-DA and sPLS-DA on a microarray data set. The PLS-DA and sPLS-DA methods are described in the Supporting Information.

#### Data description

The study investigates Small Round Blue Cell Tumors (SRBCT, Khan et al. (2001)) of 63 tumour samples with the expression levels of 2,308 genes. Samples are classified into four classes: 8 Burkitt Lymphoma (BL), 23 Ewing Sarcoma (EWS), 12 neuroblastoma (NB), and 20 rhabdomyosarcoma (RMS).

#### Unsupervised and supervised analyses

Principal Component Analysis was first applied to assess similarities between tumour types (Fig. 3**A1**). This preliminary unsupervised analysis showed no separation between tumour types, but allows to visualise the more important sources of variation, which are summarised in the first two principal components (Fig. 3**A2**). A supervised analysis with PLS-DA focuses on the discrimination of the four tumour types (Fig. 3**B1**), and led to a good performance (Fig. 3**B2**, performance assessed when adding one component at a time in the model). We then applied sPLS-DA to identify specific discriminant genes for the four tumour types. The tuning process (see ‘Choice of parameters’ Section and Electronic Supporting) led to a sPLS-DA model with 3 components and a molecular signature composed of 8, 290 and 30 genes selected on the first three components, respectively.

**Figure 3:**
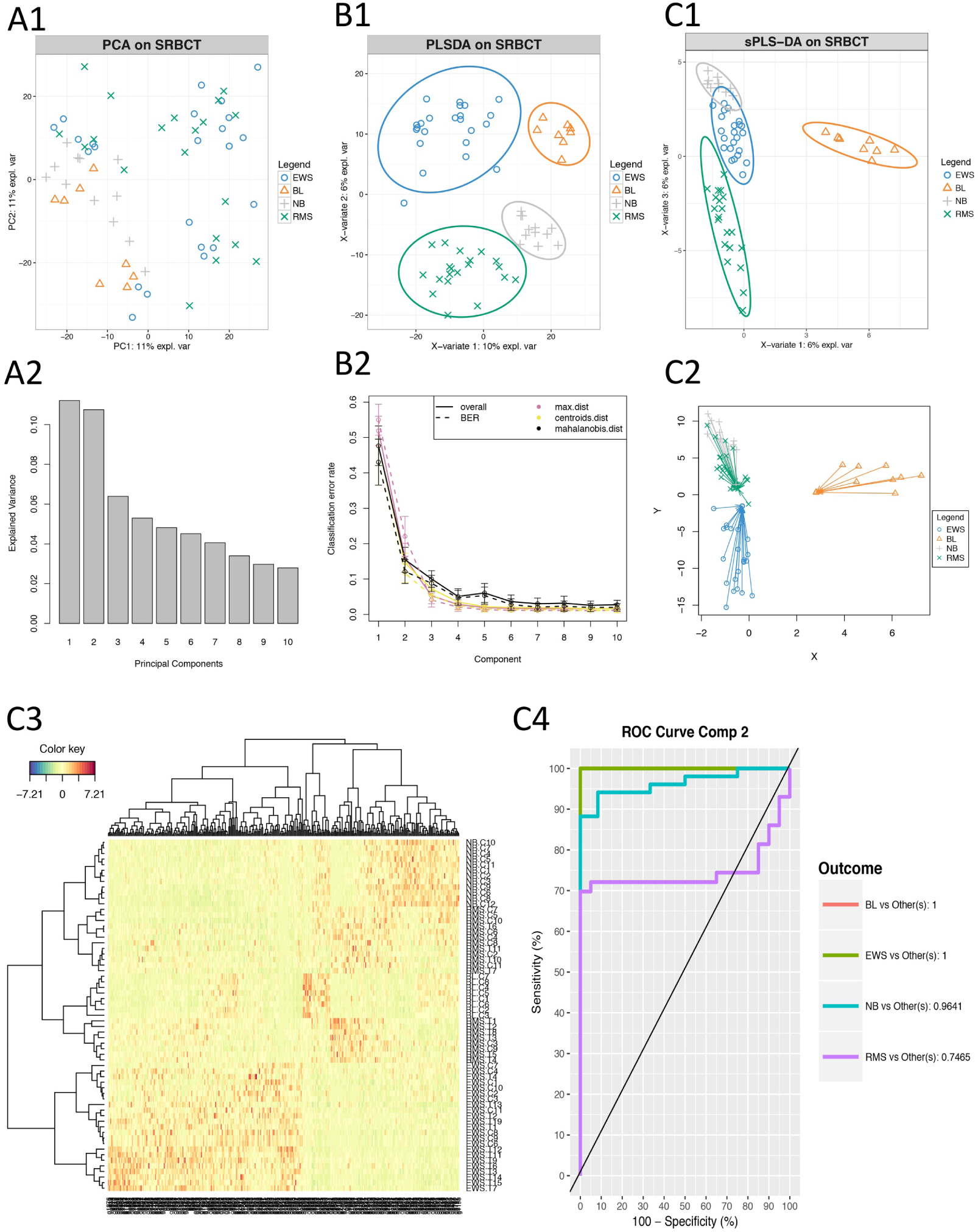
Illustration of a single ‘omics analysis with mixOmics. ***A) Unsupervised preliminary analysis with PCA***, ***A1****: PCA sample plot,* ***A2****: percentage of explained variance per component.* ***B) Supervised analysis with PLS-DA***, ***B1****: PLS-DA sample plot with confidence ellipse plots,* ***B2****: classification performance per component (overall and BER) for three prediction distances using repeated stratified cross-validation (10 × 5-fold CV).* ***C) Supervised analysis and feature selection with sparse PLS-DA***, ***C1****: sPLS-DA sample plot with confidence ellipse plots,* ***C2****: arrow plot representing each sample pointing towards its outcome category, see more details in Supporting Information*. ***C3****: Clustered Image Map (Euclidi*1*an*9 *Distance, Complete linkage) where samples are represented in rows and selected features in columns (8, 290 and 30 genes selected on each component respectively),* ***C4****: ROC curve and AUC averaged using one-vs-all comparisons.*

#### Results visualisation

The first sPLS-DA component discriminated BL vs the other tumour types (Fig. 3**C1**). The 8 genes selected on this component all had positive weight in the linear combination, and were highly expressed in BL. The second component further discriminated EWS based on 290 selected genes. The genes with a negative weight were highly expressed in EWS while the genes with a positive weight were highly expressed in either NB or RMS. Finally, the third component discriminated both NB and RMS (see Electronic Supporting). The arrow plot displays the relationship between the samples summarised as a combination of selected genes (start of the arrow) and the categorical outcome (tip of the arrow, Fig. 3**C2**).

A clustering analysis using a heatmap based on the genes selected on the first three components highlighted clusters corresponding to the four tumour types (Fig 3**C3**). ROC curve and AUC of the final model were also calculated using one-vs-all comparisons and led to satisfactory results on the first two components (Fig 3**C4**). The AUC for the first three components was 1 for all groups. Note that ROC and AUC are additional measures that may not reflect the performance of a mixOmics multivariate approaches since our prediction strategy is based on distances (see ‘Performance assessment’ Section).

#### Summary

We illustrated the mixOmics framework for the supervised analysis of a single ‘omics data set - here a microarray experiment. The full pipeline, results interpretation, associated R and Sweave codes are available in Electronic Supporting. Such an analysis suggests novel biological hypotheses to be further validated in the laboratory, when one is seeking for a *signature* of a subset of features to explain, discriminate or predict a categorical outcome. The methods has been applied and validated in several biological and biomedical studies, including ours in proteomics and microbiome Shah et al. (2016); Lê Cao et al. (2016).

### *N*-integration across multiple ‘omics data sets with DIABLO

*N*-integration consists in integrating different types of ‘omics data measured on the same *N* biological samples. In a supervised context, DIABLO performs *N*-integration by identifying a multi‘omics signature that discriminates the outcome of interest. Contrary to the concatenation and the ensemble approaches that also perform *N*-integration, DIABLO identifies a signature composed of highly correlated features across the different types of ‘omics, by modelling relationships between the ‘omics data sets Singh et al. (2016). The DIABLO method is fully described in the Supporting Information. We illustrate one analysis on a multi-‘omics breast cancer study available from the package.

#### Data description

The multi-‘omics breast cancer study includes 150 samples from three types of ‘omics: mRNA (*P*_1_ = 200), miRNA (*P*_2_ = 184) and proteomics (*P*_3_ = 142) data. Prior to the analysis with `mixOmics`, the data were normalised and filtered for illustrative purpose. Samples are classified into three subgroups: 75 Luminal A, 30 Her2 and 45 Basal.

#### Choice of parameters and analysis

As we aim to discriminate three breast cancer subtypes we chose a model with 2 components. The tuning process with the constraint model (see ‘Choice of parameters for supervised analyses’ Section and Electronic Supporting) identified a multi-‘omics signature of 5 and 8 mRNA features, 7 and 7 miRNA features and 6 and 7 proteomics features on the first two components, respectively. Sample plots of the final DIABLO model in Figure 4**A** displayed a better discrimination of breast cancer subgroups with the mRNA and proteomics data than with the miRNA data. Fig 4**B** showed that the latent components of each ‘omics were highly correlated between each others, highlighting the ability of DIABLO to model a good agreement between the data sets. The breast subtypes colors show that the components are also able to discriminate the outcome of interest.

**Figure 4:**
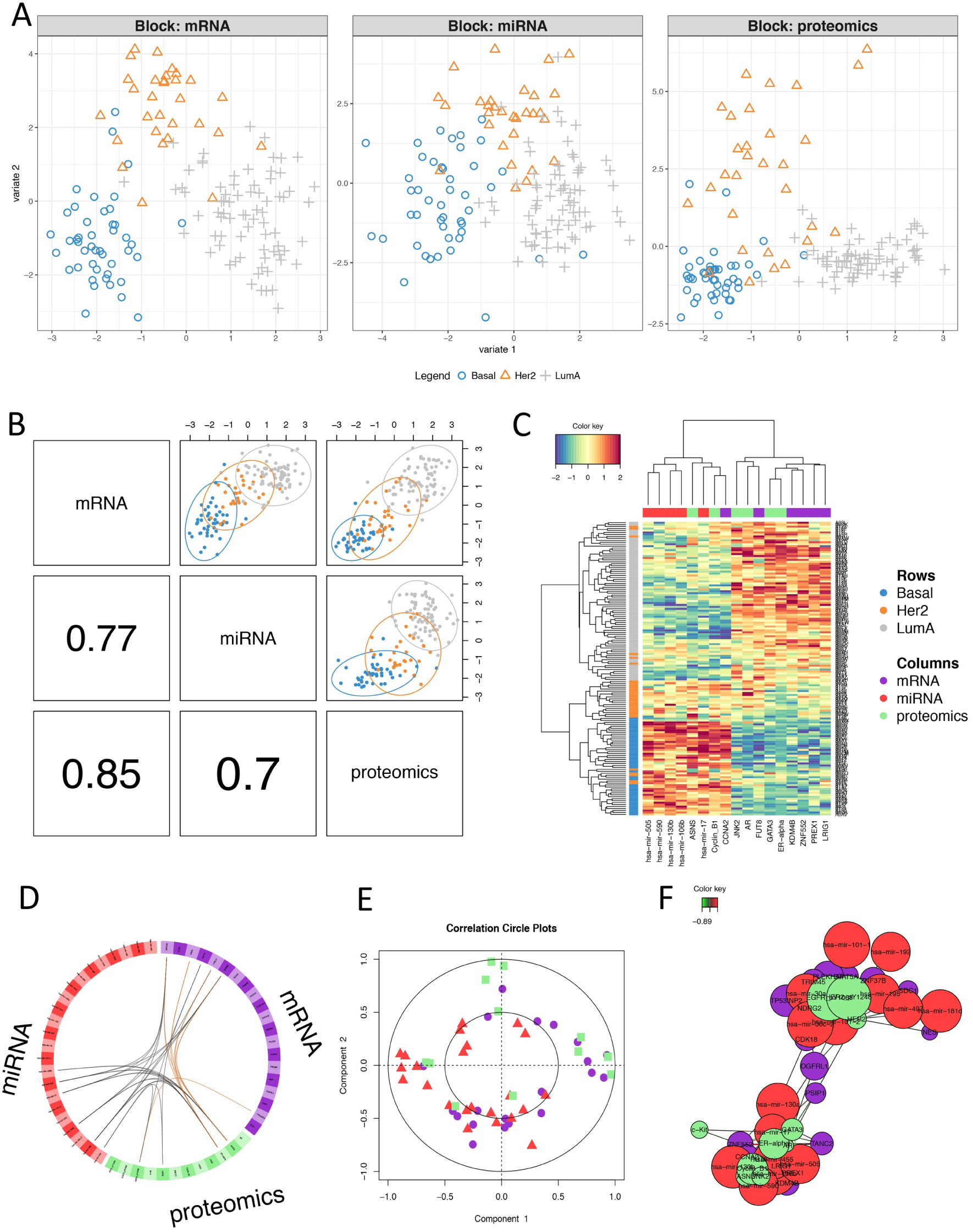
Illustration of N-integrative supervised analysis with DIABLO. ***A****: sample plot per data set,* ***B****: sample scatterplot from plotDiablo displaying the first component in each data set (upper diagonal plot) and Pearson correlation between each component (lower diagonal plot).* ***C****: Clustered Image Map (Euclidian distance, Complete linkage) of the multi-omics signature. Samples are represented in rows, selected features on the first component in columns.* ***D****: Circos plot shows the positive (negative) correlation (r >* 0.7*) between selected features as indicated by the brown (black) links, feature names appear in the quadrants,* ***E****: Correlation Circle plot representing each type of selected features,* ***F****: relevance network visualisation of the selected features*.

#### Results visualisation

Several visualisation tools are available to help the interpretation of the DIABLO results and to assess relationships between the selected multi-‘omics features (see Supporting Information and Electronic Supporting). The clustered image map (CIM) displayed a good classification of the three subtypes of breast cancer based on the 18 multi-‘omics signature identified on the first component (Fig 4**C**). The CIM output can be complemented with a circosPlot which displays the different types of selected features on a circle, with links between or within ‘omics indicating strong positive or negative correlations (Fig 4**D**). Those correlation are estimated using the latent components as a proxy, see more methodological details in González et al. (2012). We observed strong correlations between miRNA and mRNA, but only a few correlations between proteomics and the other ‘omics types. Correlation circle plots (Fig 4**E**) further highlight correlations between each selected feature and its associated latent component (see details in González et al. (2012)). The 5 miRNA features selected on the first component were highly negatively correlated with the first component (red triangles close to the (-1,0) coordinates). Contrarily, 5 of the 6 mRNA features and 4 of the 6 proteomics features selected on the first component were highly positively correlated with the first component (purple circles and green squares close to the (1,0) coordinates, respectively). Most of the features selected on the second component were close to the inner circle, which implies a weak contribution of those features to both components. Finally, a relevance network output highlighted two clusters, both including features from the three types of ‘omics (Fig 4**F**). Interactive view and .glm format are also available, see Supporting Information .

#### Summary

We illustrated the mixOmics framework for the supervised analysis of a multiple ‘omics study. The full pipeline, results interpretation and associated R and Sweave codes are available in Electronic Supporting. Our DIABLO method identifies a discriminant and highly correlated multi-‘omics signature. Predictive ability of the identified signature can be assessed (see) while the graphical visualisation tools enable a better understanding of the correlation structure of signature. Such method is the first of its kind to perform multivariate integration and discriminant analysis. DIABLO is useful to pinpoint a subset of different types of ‘omics features in those large study, posit novel hypotheses, and can be applied as a first filtering step prior to refined knowledge- and/or data-driven pathway analyses.

### *P*-integration across independent data sets with MINT

*P*-integration consists in integrating several independent studies measuring the same *P* predictors, and, in a supervised context, in identifying a robust molecular signature across multiple studies to discriminate biological conditions. The advantages of *P*-integration is to increase sample size while allowing to benchmark or compare similar studies. Contrary to usual approaches that sequentially accommodate for technical differences among the studies before classifying samples, MINT is a single step method that reduces overfitting and that predicts the class of new samples Rohart et al. (2017). The MINT method is described in Supporting Information. We illustrate the MINT analysis on a stem cell study available from the package.

#### Data description

We combined four independent transcriptomics stem cell studies measuring the expression levels of 400 genes across 125 samples (cells). Prior to the analysis with mixOmics, the data were normalised and filtered for illustrative purpose. Cells were classified into 30 Fibroblasts, 37 hESC and 58 hiPSC.

#### Choice of parameters and analysis

The optimal number of components was 1 on this data set. However, in order to obtain 2D graphics, we considered a model with 2 components. The tuning process of a MINT sPLS-DA identified a molecular signature of 6 and 16 genes on the first two components, respectively (Fig 5**A**). A MINT model based on these parameters led to a BER of 0.39 (Fig 5**B**), which was comparable to the BER of 0.37 from MINT PLS-DA when no feature selection was performed (see details in Electronic Supporting).

**Figure 5:**
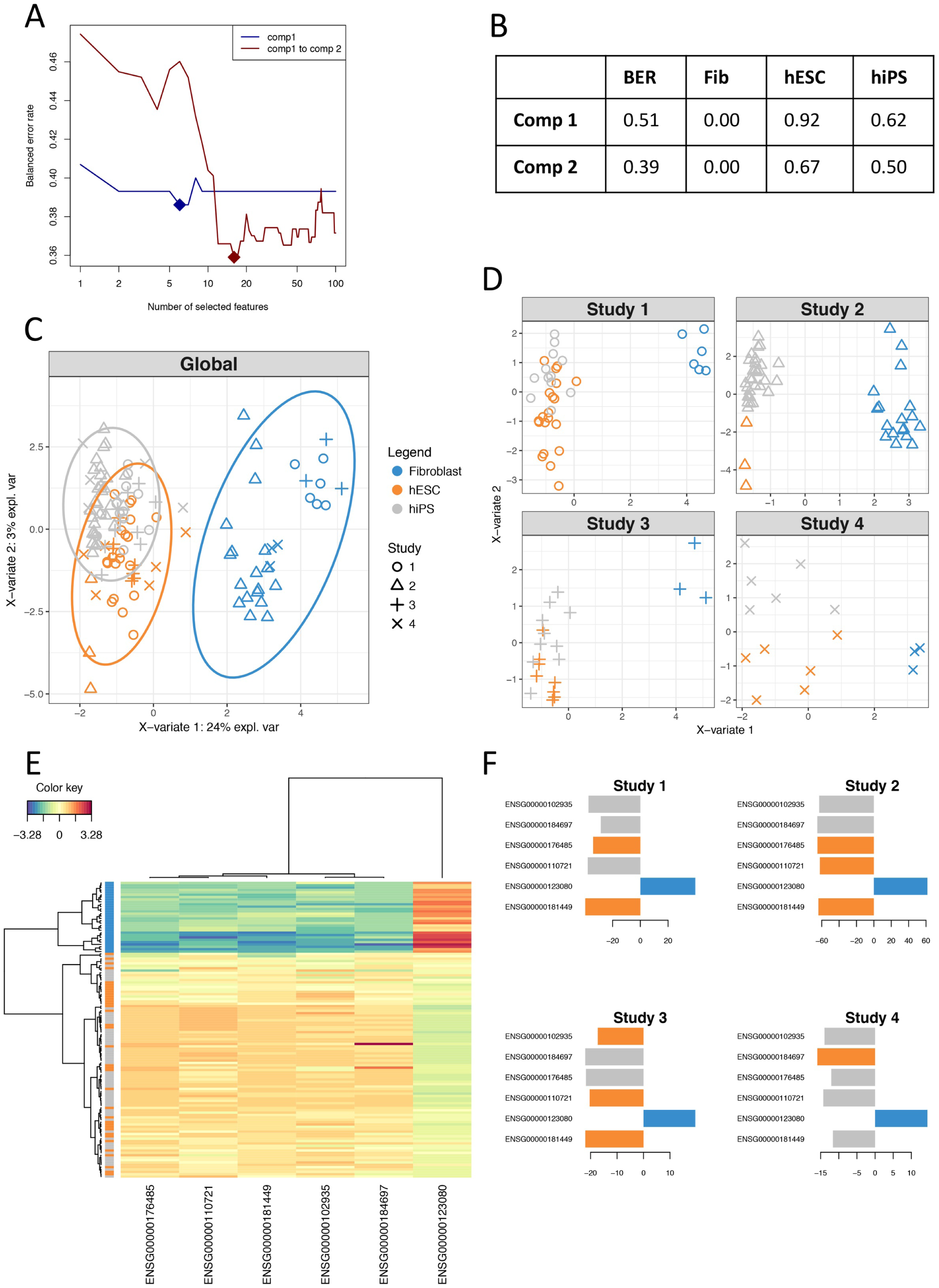
Illustration of MINT analysis in mixOmics. ***A****: Parameter tuning of a MINT sPLS-DA model with two components using Leave-One-Group-Out cross-validation and maximum distance, BER (y-axis) with respect to number of selected features (x-axis). Full diamond represents the optimal number of features to select on each component,* ***B****: Performance of the final MINT sPLS-DA model including selected features based on BER and classification error rate per class,* ***C****: Global sample plot with confidence ellipse plots,* ***D****: Study specific sample plot,* ***E****: Clustered Image Map (Euclidian Distance, Complete linkage). Samples are represented in rows, selected features on the first component in columns.* ***F****: Loading plot of each feature selected on the first component in each study, with color indicating the class with a maximal mean expression value for each gene.*

#### Results visualisation

Global sample plot (Fig 5**C**) and study-specific sample plots highlighted a good agreement between the four studies (Fig 5**D**). The first component segregated fibroblasts vs. hiPSC and hESC, and the second component hiPSC vs. hESC. Such observation was confirmed with a Clustered Image Map based on the 6 genes selected on the first component (Fig 5**E**). Importantly, the loading plots depicted in Fig 5**F** showed consistent weights assigned by the MINT model to each selected genes across each independent study.

#### Summary

We illustrated the MINT analysis for the supervised integrative analysis of multiple independent ‘omics studies. The full pipeline, results interpretation and associated R and Sweave codes are available in Electronic Supporting. Our framework proposes graphical visualisation tools to understand the identified molecular signature across all independent studies. Our own applications of the method to full data sets have showed strong potential of the method to identify reliable and robust biomarkers across independent transcriptomics studies Rohart et al. (2016, 2017).

## Conclusions and Future Directions

The technological race in high-throughput biology leads to increasingly complex biological problems which require innovative statistical and analytical tools. Our package mixOmics focuses on data exploration and data mining, which are crucial steps for a first understanding of large data sets. In this article we presented our latest methods to answer cutting-edge integrative and multivariate questions in biology.

The sparse version of our methods are particularly insightful to identify molecular signatures across those multiple data sets. Feature selection resulting from our methods help refine biological hypotheses, suggest downstream analyses including statistical inference analyses, and may propose biological experimental validations. Indeed, multivariate methods include appealing properties to mine and analyse large and complex biological data, as they allow for more relaxed assumptions about data distribution, data size and data range than univariate methods, and provide insightful visualisations. In the last few years, several R packages have been proposed for multivariate analysis as a mean for dimension reduction of one data set (see the review from Meng et al. (2016), Table 2 lists all packages and functions currently available), and the integration of two or more data sets (see Meng et al. (2016), Table 3 and factomineR Husson et al. (2017)). However, very few methods propose feature selection, including sparse CCA (pma package Witten et al. (2013)), sparse PLS (spls package, Chung et al. (2013)), penalised PLS (ppls package Kraemer and Boulesteix (2014)), sGCCA (RGCCA package Tenenhaus and Guillemot (2017)), PARAFAC and Tucker multi-way analyses (ThreeWay,PTAk, ade4 packages, Del Ferraro et al. (2015); Leibovici (2015); Thioulouse et al. (2017)) and even fewer methods provide data visualisation of the selected features (ade4).

The identification of a *combination* of discriminative features meet biological assumptions that cannot be addressed with univariate methods. Nonetheless, we believe that combining different types of statistical methods (univariate, multivariate, machine learning) is the key to answer complex biological questions. However, such questions must be well stated, in order for those exploratory integrative methods to provide meaningful results, and especially for the non trivial case of multiple data integration.

While we illustrated our different frameworks on classical ‘omics data in a supervised context, the package also include their unsupervised counterparts to investigate relationships and associations between features with no prior phenotypic or response information. Here we applied our multivariate frameworks to transcriptomics, proteomics and miRNA data. However, other types of biological data can be analysed, as well as data beyond the realm of ‘omics as long as they are expressed as *continuous values*. Sequence-based data after processing (i.e. corrected for library size and log transformed) fit this requirement, as well as clinical data. Genotype data, such as bi-allelic Single Nucleotide Polymorphism coded as counts of the minor allele can also fit in our framework, by implicitly considering an additive model. However, to consider SNPs as categorical variables additional methodological developments are required as each SNP needs to be considered as dummy indicator matrices in the sparse multivariate models.

Currently our methods are linear techniques, where each component is constructed based on a linear combination of variables. Components between different data sets however are not linearly dependent as we maximise the covariance between them Krämer and Sugiyama (2011). PLS-based models assuming a non linear relationship between different sets of data have been proposed Rosipal (2010) but the interpretation of the results in terms of identified signature is not straightforward. We are currently investigating sparse kernel-based method for non linear modelling.

Finally, the sPLS-DA framework was recently extended for microbiome 16S data Lê Cao et al. (2016), and we will further extend DIABLO and MINT for microbiome - ‘omics integration, as well as for genomic data and time-course experiments. These two promising integrative frameworks can also be combined for NP-integration, to combine multiple studies that each include several types of ‘omics data and open new avenues for large scale multiple data integration.

## Availability and requirements

The R package `mixOmics` is available from the CRAN R Core Team (2016), with tutorials and newsletter updates available from our website www.mixOmics.org.

## Availability of supporting data

The data sets supporting the results of this article are available from the mixOmics R package in a processed format. R scripts, full tutorials and reports to reproduce the results from the proposed framework are available as Sweave code from our website www.mixOmics.org.

## Author’s contributions

FR implemented the MINT method, FR, BG and AS implemented the DIABLO method, FR was the main developer of the *mixOmics* package from version 6.0.0. KALC supervises and manages the mixOmics project. FR and KALC edited the manuscript.

## Acknowledgements

FR was supported, in part, by the Australian Cancer Research Foundation (ACRF) for the Diamantina Individualised Oncology Care Centre at The University of Queensland Diamantina Institute. KALC was supported, in part, by and the National Health and Medical Research Council (NHMRC) Career Development fellowship (APP1087415). The authors would like to thank the numerous mixOmics users who continuously help in improving the usability of the package.

## Supporting Information

### S1 Text

General definitions

### S2 Text

Graphical outputs to visualise multivariate analysis results

### S3 Text

Methods description: Single ‘omics supervised multivariate analysis with PLS-DA and sPLS-DA

### S4 Text

Methods description: N-integration across multiple ‘omics data sets with DIABLO

### S5 Text

Methods description: P-integration across independent data sets with MINT

### S6 Text

Methods description: Computational time for large data sets

## Electronic Supporting Information

### E1 File

**Sweave and R codes for PLS-DA analysis** are available on our website at this link.

### E2 File

**Sweave and R codes for DIABLO analysis** are available on our website at this link.

### E3 File

**Sweave and R codes for MINT analysis** are available on our website at this link.

